# Mechanical properties of adherent cell sheets analyzed by deflection of supporting membrane

**DOI:** 10.1101/2025.04.03.647015

**Authors:** Yuri M. Efremov, Anastasia M. Subbot, Ivan A. Novikov, Sergei E. Avetisov, Peter S. Timashev

## Abstract

Multicellular structures, including cell sheets, are actively used as model systems to study intercellular interactions and can be applied in different areas of regenerative medicine. In this paper, we present a novel approach for measuring mechanical properties of cell sheets based on a simple experimental setup and numerical simulations. The advantage of the present approach is the relative ease of the sample preparation, while previous systems for the tensile tests required specialized and sensitive equipment. With the developed approach, the cell sheet on a polymer membrane is mounted in the holder on one side, and then deflection of the free end of the membrane is measured. The deflection of this cantilever-like construction depends on the elastic modulus of the membrane (which is known) and cell sheet, and also from the traction force generated by the cell sheet. By involving an experimental step with relaxing the traction force and by conducting finite element simulations, both traction force and elastic properties of the cell sheet can be estimated. We performed such measurements on cell sheets from keratocytes from corneal explants and confirmed that the developed approach is applicable for measurement of both traction force and elastic modulus of cell sheets.

## Introduction

Cell sheets have been described as two-dimensional (2D) self-organized multicellular patches (Imashiro and Shimizu, 2021; Kobayashi et al., 2019; Zurina et al., 2020), which possess intercellular interactions with paracrine signaling, accumulate ECM, and partially recreate the structural complexity of the native tissues. They were suggested to be used in regenerative medicine (Raju et al., 2020; Yamato and Okano, 2004; Zurina et al., 2023) and as a model of tissues in physiological conditions and disease to evaluate differences and enable the development of targeted therapeutics (Alghuwainem et al., 2019; Lee et al., 2018). In particular, cell sheets were used as a model of corneal diseases, keratoconus, and diabetic corneal stroma (Priyadarsini et al., 2023; Subbot et al., 2023).

While previous studies have emphasized the biological significance of cell sheets, less data is available on their mechanical properties (Efremov et al., 2021). It is important to estimate the mechanical properties of the cell sheets, for example, their contractility or elasticity, before application of such constructs in a clinical practice or in models of pathological conditions. However, estimation of mechanical parameters of cell sheets is associated with several problems. First, it is difficult to handle and attach the cell sheet in the conventional tensile test system by fixtures (clamps, pins, staples) due to the mechanical fragility of the sheet and possible damage (Uesugi et al., 2013). The cell sheet should be under conditions close to physiological during the measurements. Second, special equipment with high sensitivity is required to measure low forces during the extensional tests of soft cell sheets. Specially designed micromechanical setups were used in some works (T. Guo et al., 2023; Sorba et al., 2019). Third, the cell sheet possesses contractility due to a tension generated by the cell cytoskeleton machinery, which is an important mechanical parameter and should be separated from the elastic modulus (Heer and Martin, 2017; Stamenović and Smith, 2020).

Traditionally, tensile tests are used to analyze the mechanical properties of cell sheets (Backman et al., 2017; Harris et al., 2013; Uesugi et al., 2013). The method requires a firm attachment of the strip-shaped sample, which might be problematic due to cell sheet fragility (Uesugi et al., 2013). In particular, specially designed fixtures (Harris et al., 2013; Uesugi et al., 2013) or glue (Backman et al., 2017) were used for securing the sample edges. Alternatively, the mechanical properties of the cell sheet might be assessed using a soft elastomer membrane that also serves as a support for the cell sheet (Dassow et al., 2013; Holley et al., 2017; Sorba et al., 2019). Measuring the membrane’s mechanical behavior with and without cells allows estimation of the sheet mechanical properties. The membrane needs to be as soft as cells for accurate measurements, and a device capable of performing tensile measurements on such scales is required. The current approaches mostly do not distinguish between the elastic modulus and traction force of the cell sheet and calculate only one of the parameters.

Here, we suggested a novel approach for measuring the mechanical properties of cell sheets on supporting membranes based on a simple experimental setup and numerical simulations. Cultures of corneal fibroblasts grown in the form of cell sheets that model the corneal stroma were used. The sample stripe was clamped on one side, and then deflection of the free end of the membrane was measured. The deflection depends on the elastic modulus of the membrane (which is measured in advance) and of the cell sheet, as well as from the traction forces generated by the cell sheet. By relaxing the traction force with a specific inhibitor, the contribution of the traction force can be estimated, and then the elastic modulus can be found.

## Materials and Methods

### Cells and cell sheet preparation

Cell cultures were obtained by germinating keratocytes from corneal explants, which are fragments of the peripheral part of the cadaveric corneal discs that were not required during keratoplasty and were subject to disposal (the study was carried out in accordance with the principles of the Declaration of Helsinki, 2013). Cells of the third passage were seeded at a concentration of 300 thousand/cm^2^ in 6-well plates on the inserts with hanging permeable polycarbonate membrane, 24 mm diameter, 0.40 µm pore size (SPL, Korea). The cells were cultured for 15 days at a temperature of 37.0°C in a humid atmosphere with 5.0% CO_2_ in DMEM high glucose medium (Gibco, USA) with 2 mM glutamine (Gibco, USA), 10% fetal bovine serum (Sigma, USA), and 50 μg/ml ascorbic acid (Sigma, USA). The cell medium was changed every 2-3 days.

### Histology

To make histological sections, membranes with cells were fixed in a 10% solution of glutaraldehyde (AppliChem, Germany), embedded in Epoxy Embedding Medium (Sigma, USA), after which semi-thin sections were made at LKB Bromma Ultramicrotome System 2128 (LKB, Austria) and stained with methylene blue (Sigma, USA) and fuchsin (Sigma, USA). The imaging was performed with a “Leica DM2500” (Leica Microsystems CMS GmbH, Germany) with a camera “Leica DFC320” (Leica Microsystems CMS GmbH, Germany) in the program “Leica Application Suite”.

### Experimental setup for image acquisition of the membrane with cell sheet

To perform the mechanical analysis, the membrane with the cell layer was separated from the walls of the insert by a circular incision. Then, a strip with dimensions of 20 mm by 5 mm was cut out of the resulting circle by a single cut with a double-blade surgical scissors to avoid irregularities of the edges. No visual detachment of the cell sheet from the membrane was observed at this step.

Mechanical properties of the empty polymeric membranes were tested using the Mach-1 v500csst Micromechanical Testing System (Biomomentum Inc., Laval, Canada). The tensile strength, elongation at break, and elasticity modulus (Young’s modulus) were measured during uniaxial tension of at least three stripes (13×5 mm gauge dimensions; the thickness of 15 µm was taken from the microscopy images of the membrane cross-section) and reported as mean ± standard deviation values.

One of the short edges of the resulting strip was clamped into a precision vise (WITTE art. no. 99417, Witte GmbH, Germany) to a depth of 2 mm, and the entire structure was placed in a plastic container filled with Hanks’ Balanced Salt solution (HBSS, PanEco, Russia). The photofixation of the membrane-sheet construct was carried out with a Canon D300 camera. The relative position of the experimental camera and the camera was fixed. The scale of the images was calibrated by the known dimensions of the used clamps. Images were then analyzed by manually tracing the membrane and determining the deflection of the free end.

During the experiment with the removal of cellular traction force, the images were recorded, a myosin II inhibitor blebbistatin (Sigma-Aldrich, USA) was added to the HBSS at a concentration of 10 uM for 30 min, and then the images were recorded again. Three samples of the empty membrane and four samples with cell sheets and blebbistatin treatment were imaged in the setup in total. Paired t-test was used to compare the membrane deflection values between treated and untreated groups.

### Finite element analysis

Finite element analysis (FEA) using FEBio Studio 2.8.1 (Maas et al., 2012) was performed to investigate the behavior of cell sheets on rectangular fragments of the membrane attached to the support at one end. The load and boundary conditions were applied according to the actual experimental setup. A 2D-planar shell model was created with dimensions of 20 mm length, 5 mm width, and 15 µm thickness for the membrane, and the attached cell layer had the same length and width and 35 µm thickness. A tie constraint was employed between the membrane and cell layers. Both layers were meshed with the 20-node quadratic hexahedral elements Hex20 (200 elements, 2408 nodes). The gravity load was applied to the whole model, and the density of the materials was compensated to account for the sum of gravity and buoyancy loads. The surface traction was applied to the top of the membrane layer, thus, at the interface between cells and membrane. The elastic modulus of the membrane was selected based on the conducted mechanical measurements and was 360 MPa; the density of the membrane material was then adjusted to provide deflection of the membrane to the same value as in the experiment with the membrane in HBSS (1.01 g/cm^3^). The elastic modulus of the cell layer and the surface traction force were varied in the parametric study. In total, 64 simulations were performed with the parameters in the range of 0.1–10^6^ kPa (logarithmically spaced values) for the elastic modulus and 0–800 mN/m (linearly spaced values) for the surface traction.

## Results and discussion

When growing attached to the membrane, the cell sheet contributes to the mechanics of this combined sheet-membrane structure in two ways. First, it serves as an attached elastic layer with a certain modulus that is determined by both cellular (soft) and extracellular (ECM, generally stiffer) components. Second, the cells generate traction force on the top surface of the membrane. The cells, tissues, and organs are known to maintain a certain level of mechanical tension (stress) generated endogenously by the cytoskeletal machinery. In the attached state, this tension translates to the surface traction of the support (Tamayo et al., 2012). Therefore, both traction and elasticity of the cell sheet will affect the properties of the sheet-membrane structure. These properties were assessed here by analyzing the deflection of the cantilever-like stripe of the membrane with the attached cell sheet clamped on one side in the physiological HBSS buffer. The scheme of the experimental setup is shown in Fig. 1, and the examples of the recorded images of the deflected stripes are shown in Fig. 2.

**Figure 1.**
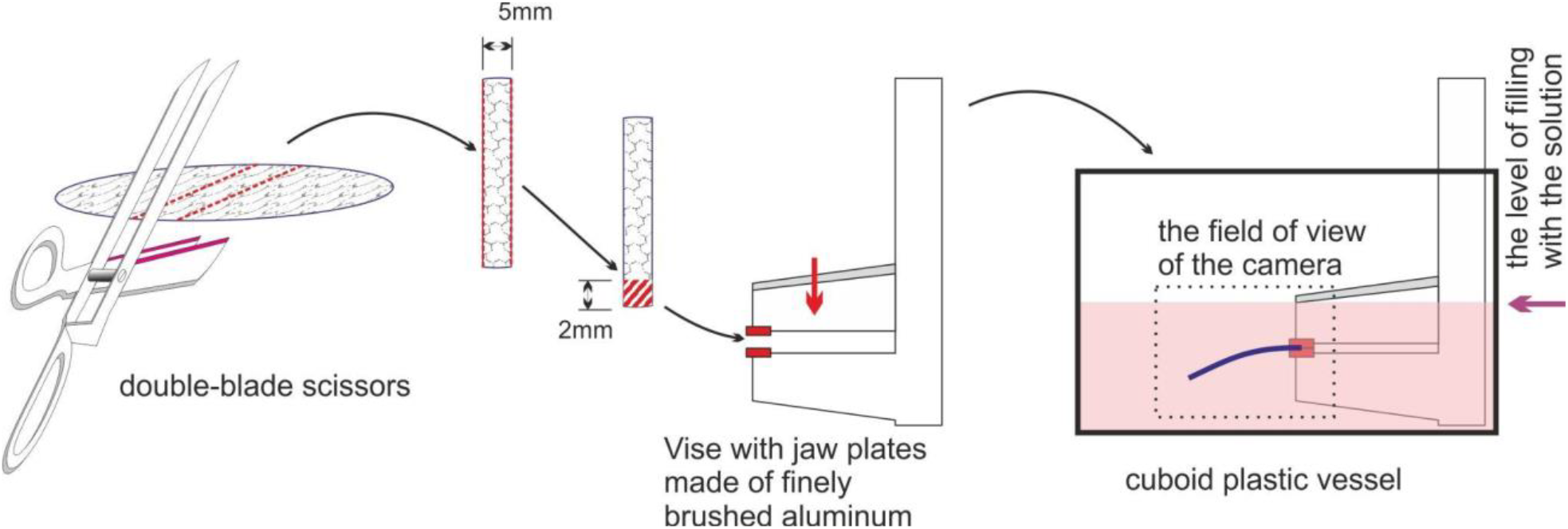
Scheme of the experimental setup for recording the deflection of the cell sheet-on-membrane strips. First, a strip (25×5 mm) was cut from the membrane by double-blade scissors. Then one end of the strip is clamped in a vice, and the vice is submerged in the vessel with HBSS.

**Figure 2.**
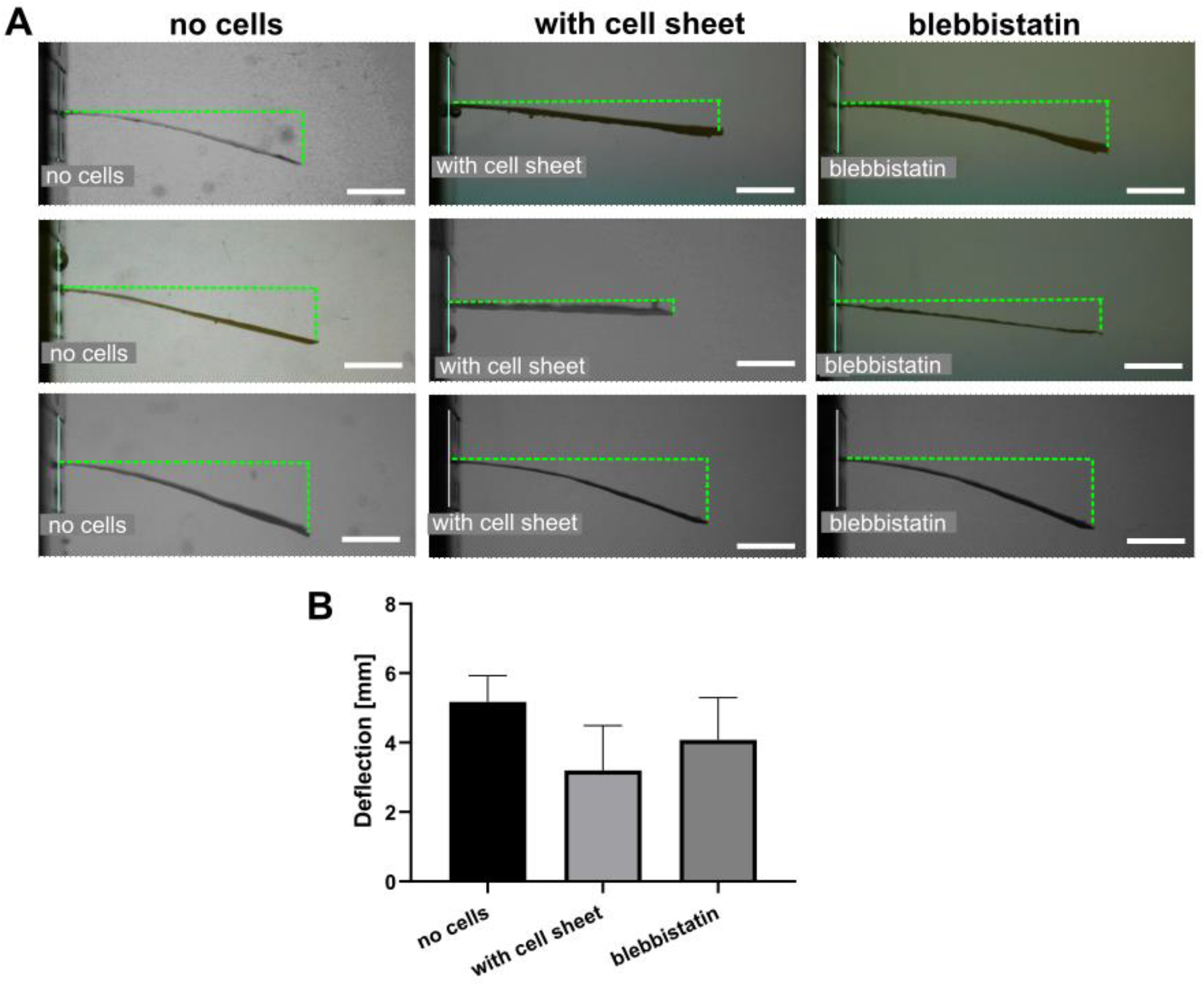
(A) Examples of the images of the membrane stripes in different conditions (all in the HBSS): empty membranes without attached cells (*no cells*); membranes with the cell sheets after 15 days of growing (*with cell sheet*); membranes with the cell sheets after incubation with blebbistatin (*blebbistatin*). Green dotted lines are shown for the assessment of the membrane deflection. Scale bars are 5 mm. (B) Measured deflection values.

Corneal keratocytes were used here for the formation of the cell sheets. They are specialized fibroblasts residing in the stroma and responsible for synthesizing its components, keeping it transparent, and repairing its wounds (West-Mays and Dwivedi, 2006; Yam et al., 2020). The cell sheets for the mechanical analysis were grown on the hanging thin polycarbonate membrane for 15 days. Histological analysis was performed to analyze the structure of the formed sheet. It confirmed the formation of the uniform multilayer structure with 25 ± 10 µm thickness (Fig. 3).

**Figure 3.**
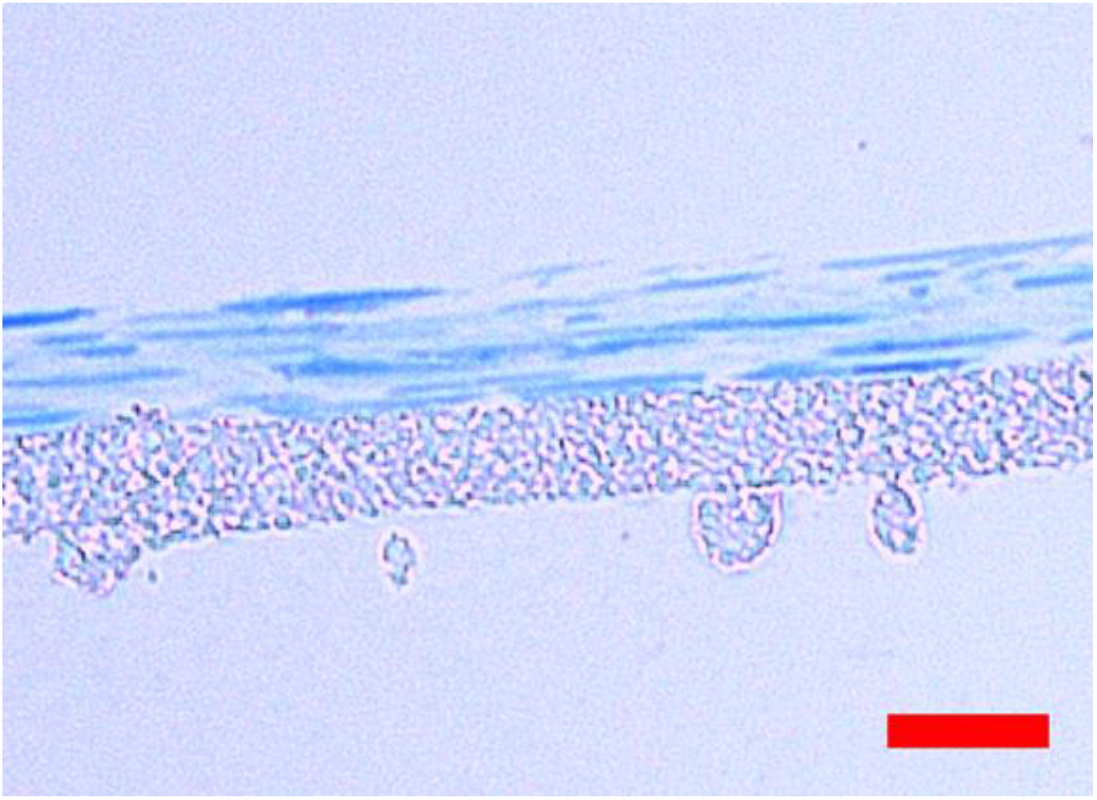
Semi-thin histological section of the cell sheet on the polymer membrane stained with methylene blue and fuchsin; the scale bar is 20 μm.

The stripes of the sheet-membrane construct were cut, clumped on one side, and side-view images were obtained in the Hanks’ Balanced Salt solution. The deflection, or shape in general, of this cantilever-like structure, as stated above, is determined by the elastic modulus and geometric sizes of the membrane and cell sheet, by the cell-generated surface traction force, and also by the density of the materials and the liquid medium. Most of the parameters can be predetermined, except the elastic modulus and traction force generated by the cell sheet; estimation of these is the purpose of the current work.

The Young’s modulus of the membrane measured in the tensional test was 360±70 MPa, while tensile strength and elongation at break were 8±1 MPa and 4±1%, respectively. The Young’s modulus value and geometrical parameters of the membrane were then taken to construct the FE model. The deflection of the membrane without the cell layer was simulated separately for the cases of the membrane in liquid and the dried membrane in air, and the results were compared with the analytical prediction of the Euler-Bernoulli beam theory for the uniformly distributed load, equation 1:

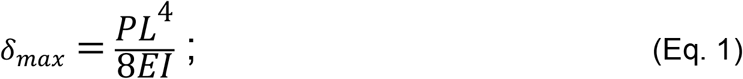

where *P* is a uniformly distributed load (a force per unit length); *L* is the length of the beam; and *E* is the elastic modulus of the material. *I* is the second moment of area of the beam’s cross-section, calculated as I = W*(H^3^)/12, where W is the width and H is the thickness of the beam. The load was estimated based on the standard gravity value, the density of the liquid medium was approximated as 1 g/mL, and the density of the membrane was estimated to be 1.01 g/mL based on the experimentally found deflection of the empty membrane in the HBSS and comparison with the theory. This value is lower than such for pure polycarbonate due to the porous structure of the membrane.

In the presence of the surface stress σ, the beam deflects, and its radius of curvature (ROC) can be estimated from Stoney’s model (Tamayo et al., 2012; Stoney, 1909):

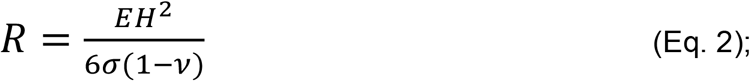

where *v* is the Poisson’s ratio of the cantilever material (assumed to be 0.45). For the small bending of the cantilever, the ROC is related to the deflection of the cantilever tip with the following equation:

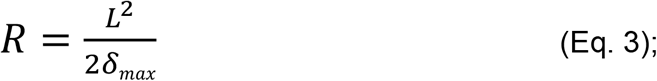

thus, the deflection can be found as:

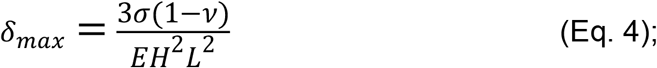

As it can be noticed, both the gravity load and surface traction cause some deflection, which is described by Equations 1 and 4, respectively.

First, the model of the membrane strip without the cell sheet was tested. In the parametric analysis, under the same gravity load, the Young’s modulus of the membrane was varied from 200 MPa to 9 GPa, and the maximum deflection was analyzed and compared with theoretical predictions. At high moduli and small deflections (below 1 mm, or 5% of the strip length), the theoretical and simulation results were in close agreement (∼10% difference). However, at larger deflections, the difference increased significantly (Fig. S1A), reaching up to ∼30% at a deflection of 8 mm (40% of the strip length). Such deviation is expected, as the Euler-Bernoulli beam theory is a simplification of the linear theory of elasticity and is valid only for small deformations.

Next, the gravity load was set to zero, and surface traction was applied and varied. Here, a deviation between the simulated results and the theoretical predictions was again observed. Although the deflection increased almost linearly with the surface traction amplitude for both theory and simulation, the slope was higher in the simulation. As a result, the deflection values were ∼10% larger in the simulation at small amplitudes (up to 1 mm), and the difference reached ∼25% at larger amplitudes (6–8 mm, or 40% of the strip length) (Fig. S1B). This discrepancy can be attributed to the fact that Stoney’s model was also developed for infinitesimally small deformations and valid only for beams that are not restrained along the edges (i.e., not clamped on one side) (10.1088/0957-4484/23/47/475702). Therefore, due to the large deflections observed in experiments, the use of finite element analysis (FEA) is more appropriate than the application of simplified theoretical models, especially since the cell layer introduces additional complexity to the system.

The FE model of the system consisting of the cell sheet on the membrane strip was constructed (Fig. 4A). The thickness of the cell sheet was taken from the histology data (35 μm). The elastic modulus of the cell layer and the surface traction were varied in the parametric study. In total, 64 simulations were performed with the parameters in the range of 0.1–10^6^ kPa (logarithmically spaced values) for the elastic modulus and 0–800 mN/m (linearly spaced values) for the surface traction. From the simulations, the whole beam profile was extracted and compared with the experimental data. The maximum deflection (deflection at the free end) of the membrane was taken to describe its overall shape and compare the simulated data with the experiment. Examples of the simulated profiles are shown in Fig 4B. It should be noted, that we selected here the representation of beam and cells as uniform homogeneous isotropic linear elastic materials as the simplified mechanical model. Besides the linear elastic model, Neo-Hookean mechanical model (Bonet and Wood, 2008) was also tested and prescribed to the membrane, cell layer, or both, but no significant differences were found in the membrane profiles under gravity load and traction (Fig. S1).

**Figure 4.**
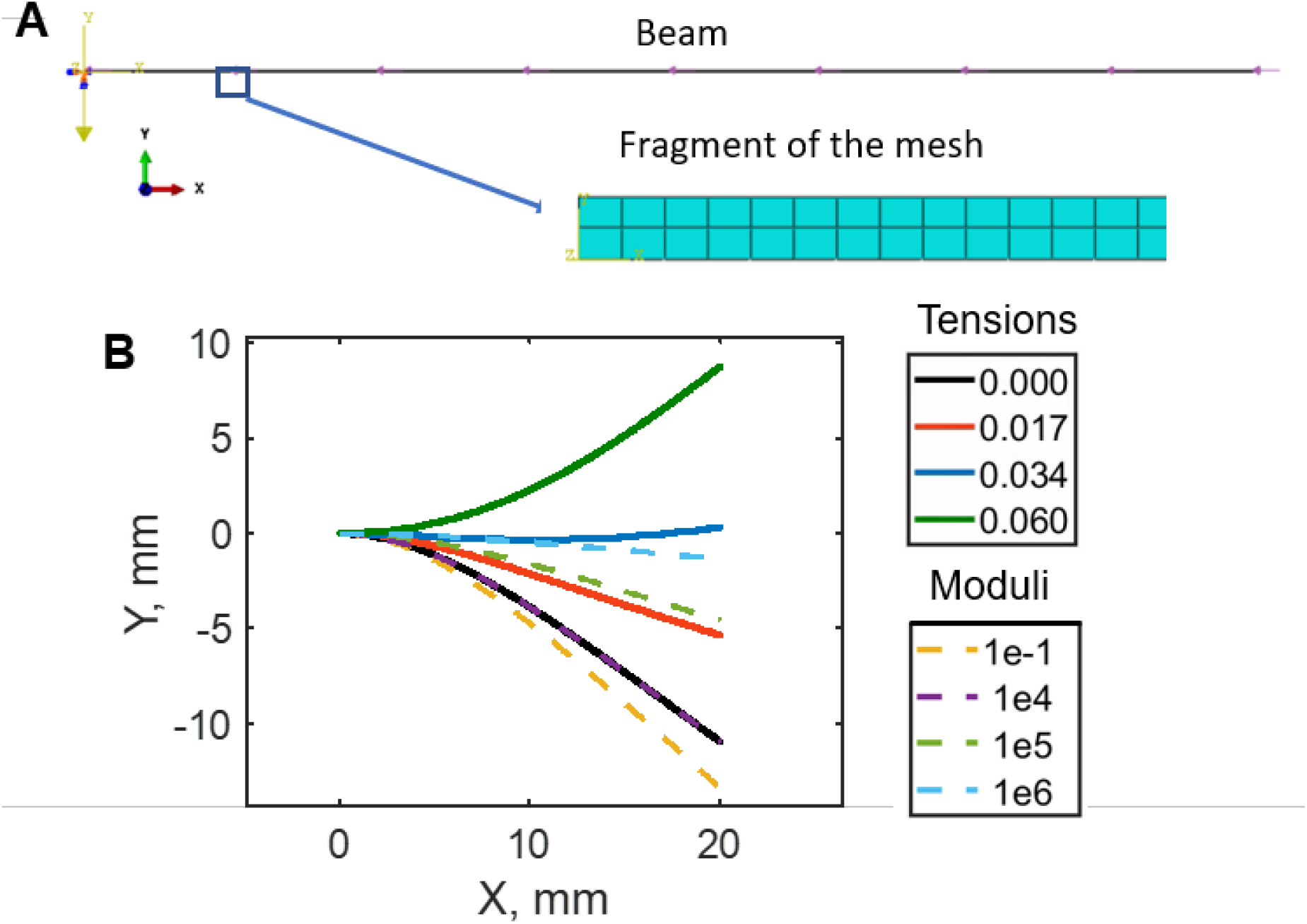
(A) 2D FE model of the cell sheet on a fragment of the membrane attached to the support at one end. Zoomed section shows a fragment of the mesh. (B) Examples of membrane profiles at different values of cell layer surface traction (at Young’s moduli of 10^4^ kPa) and Young’s moduli (at zero surface traction).

A parametric study revealed that both the elastic modulus of the cell layer and the surface traction affect the deflection of the sheet-membrane construct. The surface plot of the membrane deflection (negative values are downward deflection, positive values are upward) versus cell layer Young’s modulus and surface traction are shown in Fig. 5(A), and some particular cases are shown in Fig. 5(B, C). Since the membrane and the cell layer had comparable thicknesses, the Young’s modulus of the cell layer altered overall deflection only when it was close to or higher than the modulus of the membrane (360 MPa). The surface traction (tension), on the other hand, greatly reduced the overall deflection, affected it linearly, and even caused an upward deflection at high values. The effect of the surface traction was diminished with an increase in the cell layer’s modulus.

**Figure 5.**
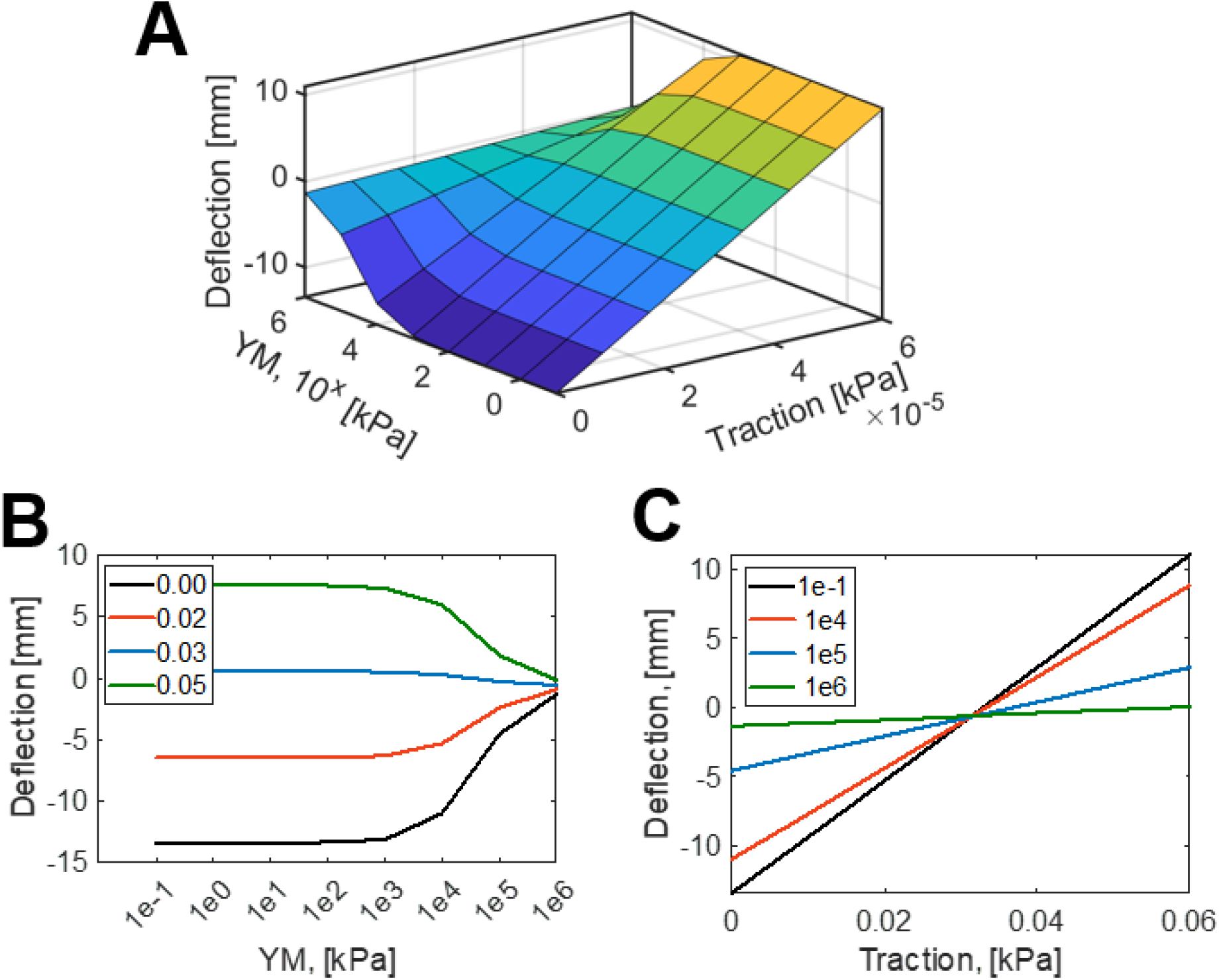
FE analysis of the deflection of the membrane with cell sheet. (A) The surface plot of the membrane deflection (negative values are downward deflection, positive – upward) versus cell layer Young’s modulus (YM, 10^x^ kPa) and surface traction (Traction, kPa, linear scale). (B, C) The plots of the membrane deflection (negative values are downward deflection, positive values are upward) versus (B) cell layer Young’s modulus at different surface tractions (see legend) and (C) surface traction at different Young’s moduli (see legend).

As can be seen from the FE analysis, both surface traction and the elastic modulus of the cell layer reduce the downward deflection of the membrane. One way to calculate both of these components is to remove the surface traction by the addition of specific cell tension inhibitors (Pietuch et al., 2013). Blebbistatin, a myosin-II ATPase inhibitor, was used here at a concentration of 10 µM. After the addition of the blebbistatin, a statistically significant increase in the downward deflection (from 3.2±1.1 mm to 4.1±1.0 mm, p=0.0367) was observed due to removal of the traction forces. Then, the deflections of the membrane with the “relaxed” cell layer (4.1±1.0 mm) and the deflection of the membrane without cells (5.2±0.6 mm) were used to estimate the elastic modulus of the cell layer. Next, given the linear dependence of the deflection from the surface traction at the certain cell layer modulus, the surface traction was estimated. In the conducted experiments, the estimated modulus of the cell layer in the relaxed state was 30±17 MPa and the surface traction was 80±40 mN/m.

The obtained parameters are close to the ones reported in the literature. The elastic modulus of the cell sheets varies in a wide range from tens of kPa to several hundred MPa depending on the cell source, preparation technique, and cultivation time (Efremov et al., 2021). High stiffness is generally assumed to be linked with a high ECM content in the sheet (Efremov et al., 2021; B. Guo et al., 2023). For comparison, the elastic modulus of the cornea was reported to be in the range of several MPa as well (Xue et al., 2018), and generation of large amounts of ECM by the used here stromal keratocytes can be expected (West-Mays and Dwivedi, 2006; Yam et al., 2020). However, we cannot exclude the possibility that blebbistatin does not remove cell traction completely, and then the elastic modulus of the cell sheet might be overestimated.

The estimated value of the traction force of a hundred mN/m is above the reported values for single cells but close to the values reported for the cell sheets (Holley et al., 2017). As reviewed in the same study (Holley et al., 2017), when converted to the traction force, values generated by single cells are generally in the range of 1-50 mN/m. It is possible that, due to the multilayered structure of cell sheet, intracellular adhesion, tension between adjacent cells, as well as cell-ECM interactions (Zuidema et al., 2020), the cell sheet can generate larger traction forces than single cells.

The developed approach has several limitations. First, viscoelastic behavior, which is often relevant for biological materials (Huang et al., 2019; Sasaki, 2012), is not included in the model. However, a stationary position of the membrane after equilibration time was analyzed, and at this point no substantial dynamic movement of the membrane was registered. Thus, the measured modulus corresponds to the long-term Young’s modulus. Second, the stiffness and tension were shown to be linked in single cells and by inhibiting tension a decrease in stiffness was observed (Martens and Radmacher, 2008; Wang et al., 2002). This linkage is much less studied in 3D cell systems, although a similar effect was shown on isolated mouse aorta as decrease in agonist-induced aortic stress and stiffness (Singh et al., 2021). Here, the measured stiffness was substantially larger than typical modulus of cells (several kPa), thus we expect larger contribution of ECM and lesser effect of cell softening on the relaxed modulus of the cell sheet. Third, the accuracy of the technique depends on the modulus of the used membrane and can be estimated to be within 1–5% of the membrane modulus (∼50 kPa for the membrane with a modulus of 360 MPa used in this study). However, by adjusting the modulus of the membrane, e.g., by using softer material such as silicone, the softer cell sheets can be analyzed.

In this study, we demonstrated an affordable and simple method to characterize both the elastic modulus and the traction force of a cell sheet. Although the setup with a cell sheet attached to the deformable membrane was used in previous studies (Dassow et al., 2013; Holley et al., 2017; Sorba et al., 2019), it generally involved the design of special experimental setups for the mechanical measurements and preparation of specific membranes, while here a commercial membrane was used that serves as a standard way to generate cell sheets. Besides that, in these works only the elastic modulus or the traction force were measured, but not both of the values simultaneously.

## Conclusions

From the mechanical point of view, cell sheets possess both stiffness (Young’s modulus) and contractility (traction force). The measurement methods that can distinguish both properties are required for the proper mechanical characterization of cell sheets. In this study, we have demonstrated an approach for measuring the elastic modulus and traction force in the cell sheets that model the corneal stroma. The benefits of the method are a relatively simple experimental setup, usage of the membranes both for the cell sheet growth and further mechanical analysis, and the possibility to use the same samples for further histology or immunohistochemistry analysis. We expect that this study will allow further development of cell sheet engineering and help in the mechanical analysis of the modeled tissue pathologies.

## Acknowledgments

This work was supported by the Russian Science Foundation (grant No. 23-74-10113, https://rscf.ru/en/project/23-74-10113/).

## References

Alghuwainem, A., Alshareeda, A.T., Alsowayan, B., 2019. Scaffold-Free 3-D Cell Sheet Technique Bridges the Gap between 2-D Cell Culture and Animal Models. IJMS 20, 4926. 10.3390/ijms20194926

Backman, D.E., LeSavage, B.L., Wong, J.Y., 2017. Versatile and inexpensive Hall-Effect force sensor for mechanical characterization of soft biological materials. Journal of Biomechanics 51, 118–122. 10.1016/j.jbiomech.2016.11.065

Bonet, J., Wood, R.D., 2008. Nonlinear Continuum Mechanics for Finite Element Analysis, 2nd ed. Cambridge University Press. 10.1017/CBO9780511755446

Dassow, C., Armbruster, C., Friedrich, C., Smudde, E., Guttmann, J., Schumann, S., 2013. A method to measure mechanical properties of pulmonary epithelial cell layers. Journal of Biomedical Materials Research - Part B Applied Biomaterials 101, 1164–1171. 10.1002/jbm.b.32926

Efremov, Y.M., Zurina, I.M., Presniakova, V.S., Kosheleva, N.V., Butnaru, D.V., Svistunov, A.A., Rochev, Y.A., Timashev, P.S., 2021. Mechanical properties of cell sheets and spheroids: the link between single cells and complex tissues. Biophysical Reviews 13, 541–561. 10.1007/s12551-021-00821-w

Guo, B., Duan, Y., Li, Z., Tian, Y., Cheng, X., Liang, C., Liu, W., An, B., Wei, W., Gao, T., Liu, S., Zhao, X., Niu, S., Wang, C., Wang, Y., Wang, L., Feng, G., Li, W., Hao, J., Gu, Q., Zhou, Q., Wu, J., 2023. High-Strength Cell Sheets and Vigorous Hydrogels from Mesenchymal Stem Cells Derived from Human Embryonic Stem Cells. ACS Appl. Mater. Interfaces 15, 27586–27599. 10.1021/acsami.3c03117

Guo, T., Zou, X., Sundar, S., Jia, X., Dhong, C., 2023. In situ measurement of viscoelastic properties of cellular monolayers via graphene strain sensing of elastohydrodynamic phenomena. Lab Chip 23, 4067–4078. 10.1039/D3LC00457K

Harris, A.R., Bellis, J., Khalilgharibi, N., Wyatt, T., Baum, B., Kabla, A.J., Charras, G.T., 2013. Generating suspended cell monolayers for mechanobiological studies. Nature Protocols 8, 2516–2530. 10.1038/nprot.2013.151

Heer, N.C., Martin, A.C., 2017. Tension, contraction and tissue morphogenesis. Development 144, 4249–4260. 10.1242/dev.151282

Holley, M.T., YekrangSafakar, A., Maziveyi, M., Alahari, S.K., Park, K., 2017. Measurement of cell traction force with a thin film PDMS cantilever. Biomed Microdevices 19, 97. 10.1007/s10544-017-0239-3

Huang, D., Huang, Y., Xiao, Y., Yang, X., Lin, H., Feng, G., Zhu, X., Zhang, X., 2019. Viscoelasticity in natural tissues and engineered scaffolds for tissue reconstruction. Acta Biomaterialia 97, 74–92. 10.1016/j.actbio.2019.08.013

Imashiro, C., Shimizu, T., 2021. Fundamental Technologies and Recent Advances of Cell-Sheet-Based Tissue Engineering. International Journal of Molecular Sciences 22, 425. 10.3390/ijms22010425

Kobayashi, J., Kikuchi, A., Aoyagi, T., Okano, T., 2019. Cell sheet tissue engineering: Cell sheet preparation, harvesting/manipulation, and transplantation. Journal of Biomedical Materials Research - Part A 107, 955–967. 10.1002/jbm.a.36627

Lee, J., Shin, D., Roh, J.-L., 2018. Development of an in vitro cell-sheet cancer model for chemotherapeutic screening. Theranostics 8, 3964–3973. 10.7150/thno.26439

Maas, S.A., Ellis, B.J., Ateshian, G.A., Weiss, J.A., 2012. FEBio: Finite Elements for Biomechanics. Journal of Biomechanical Engineering 134, 011005. 10.1115/1.4005694

Martens, J.C., Radmacher, M., 2008. Softening of the actin cytoskeleton by inhibition of myosin II. Pflugers Archiv European Journal of Physiology 456, 95–100. 10.1007/s00424-007-0419-8

Pietuch, A., Brückner, B.B.R., Fine, T., Mey, I., Janshoff, A., 2013. Elastic properties of cells in the context of confluent cell monolayers: impact of tension and surface area regulation. Soft Matter 9, 11490–11502. 10.1039/c3sm51610e

Priyadarsini, S., McKay, T.B., Escandon, P., Nicholas, S.E., Ma, J.-X., Karamichos, D., 2023. Cell sheet-based approach to study the diabetic corneal stroma. Experimental Eye Research 237, 109717. 10.1016/j.exer.2023.109717

Raju, R., Oshima, M., Inoue, Miho, Morita, T., Huijiao, Y., Waskitho, A., Baba, O., Inoue, Masahisa, Matsuka, Y., 2020. Three-dimensional periodontal tissue regeneration using a bone-ligament complex cell sheet. Sci Rep 10, 1656. 10.1038/s41598-020-58222-0

Sasaki, N., 2012. Viscoelastic Properties of Biological Materials, in: De Vicente, J. (Ed.), Viscoelasticity - From Theory to Biological Applications. InTech. 10.5772/49979

Singh, K., Kim, A.B., Morgan, K.G., 2021. Non-muscle myosin II regulates aortic stiffness through effects on specific focal adhesion proteins and the non-muscle cortical cytoskeleton. J Cellular Molecular Medi 25, 2471–2483. 10.1111/jcmm.16170

Sorba, F., Poulin, A., Ischer, R., Shea, H., Martin-Olmos, C., 2019. Integrated elastomer-based device for measuring the mechanics of adherent cell monolayers. Lab on a Chip 19, 2138–2146. 10.1039/c9lc00075e

Stamenovic, D., Smith, M.L., 2020. Tensional homeostasis at different length scales. Soft Matter 16, 6946–6963. 10.1039/d0sm00763c

Stoney, G.G., 1909. The tension of metallic films deposited by electrolysis. Proc. R. Soc. Lond. A 82, 172–175. 10.1098/rspa.1909.0021

Subbot, A.M., Novikov, I.A., Patejuk, L.S., Kobzeva, A.V., Avetisov, S.Je., 2023. ?ell model for experimental research in keratoconus pathogenesis. Genes & Cells 18, 69–77. 10.23868/gc321383

Tamayo, J., Ruz, J.J., Pini, V., Kosaka, P., Calleja, M., 2012. Quantification of the surface stress in microcantilever biosensors: revisiting Stoney’s equation. Nanotechnology 23, 475702. 10.1088/0957-4484/23/47/475702

Uesugi, K., Akiyama, Yoshitake, Hoshino, T., Akiyama, Yoshikatsu, Yamato, M., Okano, T., Morishima, K., 2013. Measuring mechanical properties of cell sheets by a tensile test using a self-attachable fixture. Journal of Robotics and Mechatronics 25, 603–610. 10.20965/jrm.2013.p0603

Wang, N., Tolić-Nørrelykke, I.M., Chen, J., Mijailovich, S.M., Butler, J.P., Fredberg, J.J., Stamenović, D., 2002. Cell prestress. I. Stiffness and prestress are closely associated in adherent contractile cells. American Journal of Physiology-Cell Physiology 282, C606–C616. 10.1152/ajpcell.00269.2001

West-Mays, J.A., Dwivedi, D.J., 2006. The keratocyte: Corneal stromal cell with variable repair phenotypes. The International Journal of Biochemistry & Cell Biology 38, 1625–1631. 10.1016/j.biocel.2006.03.010

Xue, C., Xiang, Y., Shen, M., Wu, D., Wang, Y., 2018. Preliminary Investigation of the Mechanical Anisotropy of the Normal Human Corneal Stroma. Journal of Ophthalmology 2018, 1–7. 10.1155/2018/5392041

Yam, G.H.F., Riau, A.K., Funderburgh, M.L., Mehta, J.S., Jhanji, V., 2020. Keratocyte biology. Experimental Eye Research 196, 108062. 10.1016/j.exer.2020.108062

Yamato, M., Okano, T., 2004. Cell sheet engineering. Materials Today 7, 42–47. 10.1016/S1369-7021(04)00234-2

Zuidema, A., Wang, W., Sonnenberg, A., 2020. Crosstalk between Cell Adhesion Complexes in Regulation of Mechanotransduction. BioEssays 42, 2000119. 10.1002/bies.202000119

Zurina, I.M., Presniakova, V.S., Butnaru, D.V., Svistunov, A.A., Timashev, P.S., Rochev, Y.A., 2020. Tissue engineering using a combined cell sheet technology and scaffolding approach. Acta Biomaterialia 113, 63–83. 10.1016/j.actbio.2020.06.016

Zurina, I.M., Presniakova, V.S., Butnaru, D.V., Timashev, P.S., Rochev, Y.A., Liang, X.-J., 2023. Towards clinical translation of the cell sheet engineering: Technological aspects. Smart Materials in Medicine 4, 146–159. 10.1016/j.smaim.2022.09.002

